# Recurrent connections enable point attractor dynamics and dimensionality reduction in a connectome-constrained model of the insect learning center

**DOI:** 10.1101/2024.01.10.574960

**Authors:** Justin Joyce, Raphael Norman-Tenazas, Patricia Rivlin, Grace M. Hwang, Isaac Western, Kechen Zhang, William Gray-Roncal, Brian Robinson

## Abstract

The learning center in the insect, the mushroom body (MB) with its predominant population of Kenyon Cells (KCs), is a widely studied model system to investigate neural processing principles, both experimentally and theoretically. While many computational models of the MB have been studied, the computational role of recurrent connectivity between KCs remains inadequately understood. Dynamical point attractors are a candidate theoretical framework where recurrent connections in a neural network can enable a discrete set of stable activation patterns. However, given that detailed, full recurrent connectivity patterns in biological neuron populations are mostly unknown, how theoretical models are substantiated by specific networks found in biology has not been clear. Leveraging the recent release of the full synapse-level connectivity of the MB in the fly, we performed a series of analyses and network model simulations to investigate the computational role of the recurrent KC connections, especially their significance in attractor dynamics. Structurally, the recurrent excitation (RE) connections are highly symmetric and balanced with feedforward input. In simulations, RE facilitates dimensionality reduction and allows a small set of self-sustaining point attractor states to emerge. To further quantify the possible range of network properties mediated by RE, we systematically explored the dynamical regimes enabled by changing recurrent connectivity strength. Finally, we establish connections between our findings and potential functional or behavioral implications. Overall, our work provides quantitative insights into the possible functional role of the recurrent excitatory connections in the MB by quantifying the point attractor network dynamics within a full synapse-level connectome-constrained highly recurrent network model. These findings advance our understanding of how biological neural networks may utilize point attractor dynamics.

**Author summary:** Point attractor neural networks are widely used theoretical models of associative memory, where recurrent connections between neurons enable a discrete set of stable activation patterns that can recover a full pattern based on partial cues. The detailed full recurrent connectivity patterns in biological neuron populations are largely unknown, however, raising questions about the precise correspondence between theoretical point attractor models and neural networks found in biology. Recent breakthroughs have unveiled the synapse-level connectivity of all neurons within the learning center of an insect, including recurrent connections between the primary neuron type—a crucial component with an elusive computational role. In this work, we perform analyses of these recurrent connectivity patterns and simulate neural network models that have these biologically constrained neural network patterns. We find that these recurrent connections are highly symmetric and balanced with input to the memory center. In simulations, we find that these recurrent connections perform dimensionality reduction and enable a small set of point attractor states. We additionally characterize how the strength of these recurrent connections affects network properties and downstream behavioral consequences. Overall, this work advances an understanding of the insect learning center as well as the relationship between theoretical and biological recurrent networks.

## Introduction

Point attractor dynamics can be used as a theoretical framework [4, 37, 27] for understanding the collective computational properties of interacting neurons in recurrent neural networks and the time-varying relationship of the emergent neural activation patterns. A prototypical theoretical example of point attractor dynamics in recurrent networks is the Hopfield network, where recurrent connections governed by a Hebbian learning rule or pairwise correlations of activities can store memory patterns that correspond to stable equilibrium states [24, 22]. This type of recurrent point attractor network has desirable emergent properties such as recall from partial cues (i.e., pattern completion), in addition to robustness against dynamical noise and structural damages. Recurrent networks can embed both discrete point attractors and continuous attractors [42] which may computationally represent discrete or continuous variables and be utilized for tasks such as abstract optimization [23], decision-making [8], and navigation [36], [28] While attractor dynamics have rich theoretical computational properties, it is still unknown to what degree biological networks utilize point attractor dynamics and how the architecture of biological point attractor networks relate to their theoretical counterparts.

Dynamical attractors in recurrent networks are increasingly used as a theoretical framework for the understanding and interpretation of neurophysiological data, including cortical data [14, 10, 26]. Attractor dynamics have been investigated for their computational role in neural regions, including representation of continuous spatial variables in the hippocampal system [36], [28], and spatial working memory in the prefrontal cortex [13, 52]. Point attractor dynamics are a substrate for the long-term storage of patterns in recurrent neural connections; CA3 in the hippocampus exhibits highly recurrent connectivity and has long been thought of as a candidate for attractor dynamics [50, 30, 41]. A critical challenge for investigation of attractor dynamics in most neural regions, however, is that the detailed connectivity of neural populations at the synapse-level is not known.

The memory center in insects is the mushroom body (MB) [20], with detailed connectivity at the synapse-level recently available [16], [32], [44], [48]. The computational functions of neurons in the MB in memory formation, including projection neurons (PNs), Kenyon cells (KCs), the anterior paired lateral neurons (APLs), and MB output neurons (MBONs), have been extensively studied theoretically and experimentally [38]. Olfactory stimuli are encoded in olfactory receptor neurons (ORNs), which synapse onto a smaller population of PNs. Sensory input from the PNs is projected onto a larger number of KCs, the most numerous neuron type in the MB. Plasticity in the connectivity between KCs and a smaller number of MBONs is sufficient to drive learning of associations of sensory signals [53]. Subsequent MBON activation by processed sensory input after learning has been shown to mediate numerous biological functions, namely the attraction or aversion to appropriate stimuli, sleep promotion, and wake promotion [5]. While multiple computational principles have been investigated in the MB [11, 38], a common principle in the computational role of the MB is the feedforward processing from PNs to KCs to MBONs with feedback inhibition (FBI) provided by APLs enabling competition between neurons and sparsification of neural activation patterns. This predominantly feedforward network has algorithmic features such as performing expansion recoding [34], pattern separation [33], and locality-sensitive hashing [15]. Missing from a more comprehensive understanding, however, are recurrent excitatory (RE) connections between KC neurons.

Here, we investigated the computational role of the RE connections in KCs in the MB by quantifying the connectivity patterns from synapse-level connectome data [44], as well as by performing a series of network simulations. In our connectivity analyses, we found consistent features of input integration and identify elevated symmetry patterns suggestive of a role in supporting point attractor dynamics. We create network simulations that process olfactory inputs from olfactory sensory patterns through MBONs, incorporating recurrent KC connections. In simulations, we demonstrate how recurrent connections change network output and achieve dimensionality reduction. Additionally, we investigate how recurrent connectivity can mediate a small set of self-sustaining point attractor states. Furthermore, we characterize attractor dynamics and network activity regimes as a function of relative recurrent excitation, inhibition, and assumptions of connectome parsing.

## Results

### Network Structure

A synapse-level connectome constrained network is created that models the signal transformation of olfactory input through the MB by utilizing synapse counts between neurons as the effective connectivity weight (Fig 1). The input to the network, 2569 ORNs, synapse with PNs (Fig 1B), which subsequently synapse onto downstream KCs (Fig 1D) in a feedforward architecture. KCs, the most numerous neural population in the MB, are organized into three major classes: *α*/*β*, *α^′^*/*β^′^*, and *γ*—named for the MB lobe that they predominantly occupy. These classes further subdivide, based on genetic driver lines, immunohistochemistry and single cell morphology, into a total of 14 subtypes. Olfactory PNs connect to all KCs, providing the majority of connections to all subtypes except two, KC*α*/*β*-p and KC*γ*-d, which receive predominantly visual inputs [32]. KCs are presynaptic, postsynaptic, or both to all neuron populations in the network model except for ORNs. Besides feedforward input from PNs, other sources of input to KCs include recurrent excitation (RE) from other KCs as well as feedback inhibition (FBI) from the APL neuron. These excitatory and inhibitory connections account for the majority of pre- and post-synaptic connections (54%) with KCs. Recurrence between KCs is the strongest within the same KC lobe (Fig 1F), wherein neurons are densely connected, ranging from 7.2% of possible neuron connections in the KC*γ*-m subtype to 50.1% of possible neuron connections in the KC*α*/*β*-p subtype.

**Fig 1.**
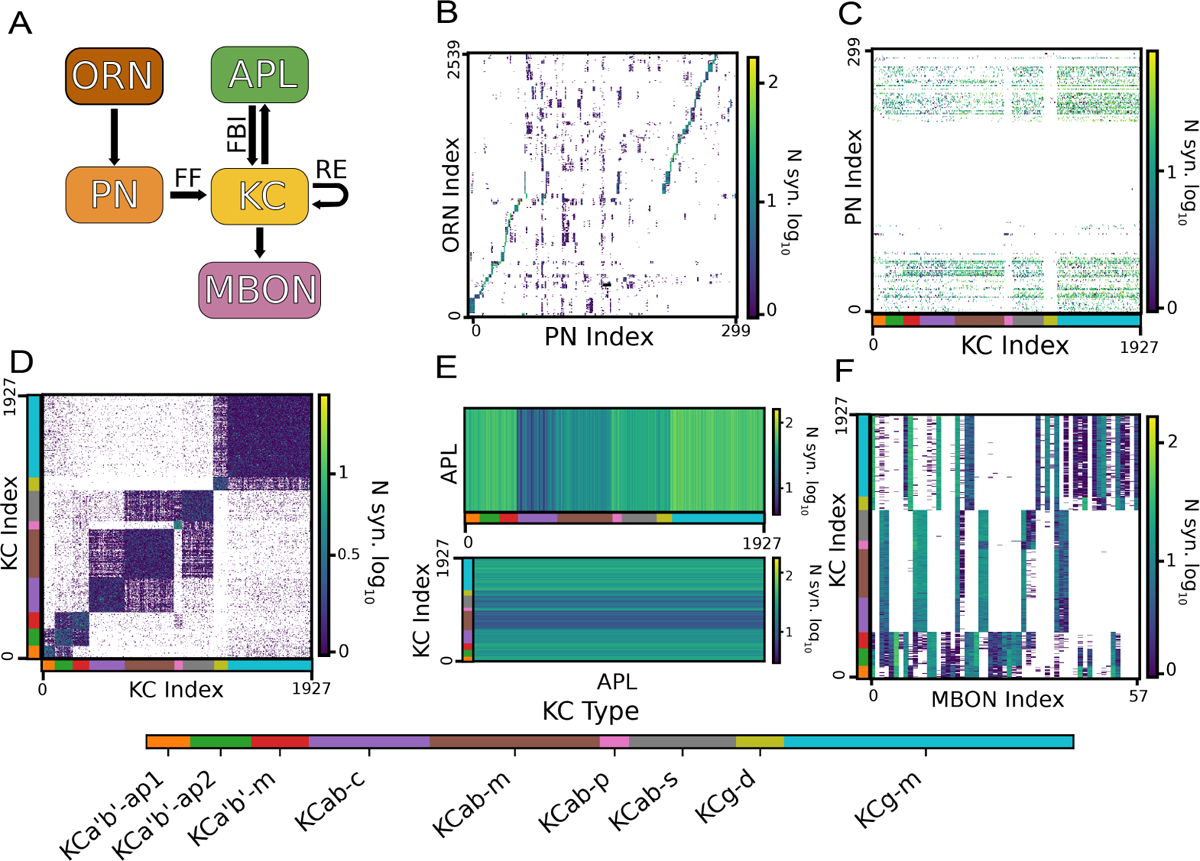
Synapse-level connectome constrained network model of olfactory processing in the MB. (A) Connectivity diagram of the modeled neuron populations, with arrows indicating the source and target neuron populations. Sources of input for the KCs include feedforward (FF) input from PNs, feedback inhibition (FBI) from the APL neuron, and recurrent excitation (RE). (B-F) Connectivity weight matrices between neuron populations were defined by queried synapse counts between neurons (white signifying no connection). Neurons were ordered alphabetically by instance name, resulting in groupings by olfactory glomerulus (ORN, PN), subtype (KC), and lobe (MBON) for the neuron populations. The subtype of KC neurons is represented by color for ease of reference, with a key at the bottom of the figure.

Comparatively, connections that cross between different subtypes or major classes are rare, ranging from 0.9% to 4.7% of possible connections in the least and most densely connected subtypes (from KC*α*’/*β*’-m to KC*α*/*β*-p and from KC*α*’/*β*’-ap2 to KC*α*’/*β*’-m, respectively). The connections from the KCs to the anterior paired lateral neuron (APL) and the corresponding connections from the APL back to the KCs form the inhibitory feedback loop, which demonstrates lobe-specific patterns of connectivity. We see that among subtypes that receive predominantly olfactory input, the *α*/*β* class (in the *α*/*β*-m and *α*/*β*-s subtypes) contributes significantly fewer connections to the APL than the *α*’/*β*’ and *γ* classes (Fig 1C). We also see that the KC*α*/*β*-c and KC*α*/*β*-m receive very few APL inputs compared to other KC neurons. The connections between KCs and MBONs are known to be compartmentalized by input lobe [5] with clear delineations between MBONS that primarily receive input from specific subtypes (Fig 1E). In sum, the synapse-level connectome constrained network model includes highly structured connectivity patterns that reflect glomeruli, subtype, and lobe-specific variability.

The properties of input integration in the network can be analyzed by quantifying the relative number of input synapses each KC receives from RE, FBI, and FF input sources. The correlation between each pair of KC inputs (FF, RE, and FBI) is calculated to quantify the relationships between the input types within each neuron. A given KC’s number of PN input synapses and recurrent KC-KC input synapses have a positive correlation of r=0.63 (Fig 2B). The number of APL input synapses, however, shows a smaller correlation of r=0.41 and 0.37 to the KC and PN input synapses respectively (2C-D). Each correlation shown includes all KC subtypes that predominantly receive sensory input from olfactory neurons; this group excludes the *α*/*β*-p and *γ*-d subtypes. This shows that KCs that have a greater number of synapses from sensory neurons are more likely to have a greater number of excitatory synapses from other KCs than inhibitory synapses from the APL. In other words, a high number of sensory synapses is more predictive of recurrent excitation than inhibition. The observed balance of feedforward sensory input with recurrent excitation is thus the most consistent trend observed in the integration of different input types for KCs.

**Fig 2.**
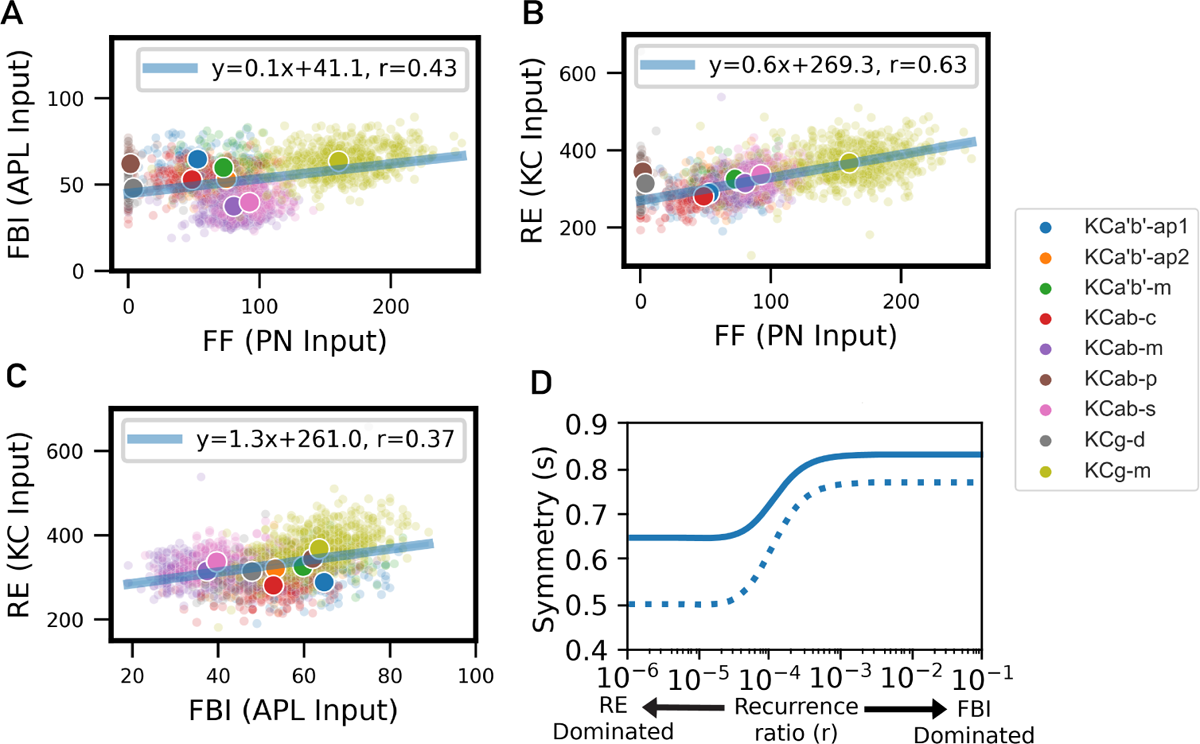
Kenyon cell connectivity structure characterization. (A-C) The balance of input types (FF, RE, and FBI, compared pairwise) per KC. The smaller markers signify the number of synapses of each input type for each KC, colored by KC subtype (larger markers signify the mean per subtype). All input types were positively correlated, with RE and FF exhibiting the highest correlation. Note that the non-olfactory KC subtypes (*α*/*β*-p and *γ*-d) were excluded from the line of best fit and correlation analysis. (D) The symmetry of the composite recurrent KC weight matrix (solid line) at all relative levels of FBI and RE (lowest *s* is RE-dominated, highest *s* is FBI-dominated) was higher than the 0.5 expected from a standard normal distribution. A comparison to a null structure baseline (dotted line) reflects that the structure of the RE component increased symmetry more than the FBI component (whereas the higher symmetry of the FBI component was driven more by its general distribution rather than its structure).

We analyze the symmetry of the recurrent connectivity between KCs to characterize the relationship between the observed connectivity and the symmetric connectivity patterns in theoretical models of point attractors. Given the existence of both recurrent excitation and feedback inhibition contributing to recurrent connectivity, a composite recurrent weight matrix is used for analysis, *W_R_* = *W_RE_ − rW_F BI_*, where *W_RE_* is the KC-KC recurrent excitation and *W_F BI_* = *W_KC→AP L_W_AP L→KC_* captures properties of feedback inhibition. The composite weights *W_R_* would be the total effective recurrent weights if the APL neuron were perfectly linear. By varying a positive-valued recurrence ratio scalar, *r*, the symmetry can be quantified over varying relative contributions of RE and FBI ranging from RE-dominated (low *r*) to FBI-dominated (high *r*). A measure of symmetry, *S*, is defined by using the Euclidean norms of the symmetric and anti-symmetric components of matrix *W_R_*:

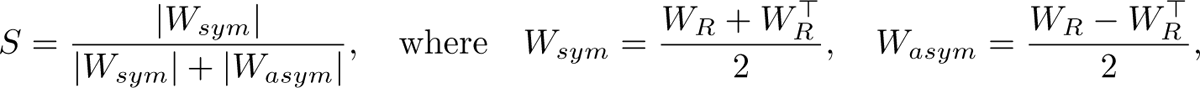

with *⊤* indicating transpose. The symmetry measure is bound between 0 and 1, with 0 indicating perfect asymmetry, 1 indicating perfect symmetry, and 0.5 indicating the symmetry expected from a weight matrix drawn from a standard normal distribution. The measured symmetry is always above 0.5 when varying levels of RE and FBI, ranging from 0.65 to 0.83 when the composite recurrent connectivity is dominated by RE and FBI, respectively. The measured symmetry is affected by both the distribution of an individual entry and the structure of the connectivity involving different entries—for example, a perfectly homogeneous distribution would have a measured symmetry value of 1. To measure the contribution of symmetry based on the structure of the recurrent connectivity (as opposed to the general distribution of the weights), the symmetry of *W_R_* is compared to a null structure baseline, created by randomly permuting *W_R_*at each level of *S*. The differences in the null structure symmetry values between the RE-dominated and FBI-dominated recurrent weights reflect the general distribution of weights, and the measured value of 0.77 in the FBI-dominated null structure weights reflects the value from a more homogeneous distribution. The structure of the connectivity from the RE-dominated composite weight matrix demonstrates a larger increase of 0.15 from the null structure baseline (0.65 vs. 0.5), relative to the 0.06 (0.83 vs. 0.77) observed in the FBI-dominated composite weight matrix. In general, the observed recurrent connectivity has symmetry at levels of above chance, with the structure of the RE showing a larger relative contribution to symmetry than FBI.

### Recurrence and Inhibition

To investigate the functional role of recurrent connectivity, we simulated network activation across samples of sensory input patterns. Full details of the network structure, network input, neuron models, and network variants are outlined in the Material and Methods section. Simulations utilize olfactory input patterns, which are resampled from experimentally observed activation patterns to olfactory stimuli cues (Fig 3A). The sampled ORN input patterns (Fig 3C) drive PN activation patterns (Fig 3E), which in turn provide the feedforward input to KCs (Fig 3G). We investigate the independent roles of FBI and RE by adding each individually with a scaled version of the FBI or RE weight matrix, which decreases or increases the KC activation patterns by 15% respectively in mean activity from the patterns produced by feedforward input alone at the final time step. With only feedforward input, neuron activation monotonically increases to a steady state pattern (Fig 3G). When inhibition is added, the overall activity of neurons is decreased, and neurons with low firing levels become inactive. In this way, FBI decreases the number of neurons active at the final state and enables competition (Fig 3D). When RE alone is added, as activity increases, RE changes the pattern of activated KCs (Fig 3F). Additionally, FF and FBI show bands of low activity in the KC subtypes KC*α*/*β*-p and KC*γ*-d that result from weak connectivity from the olfactory PNs. Contrarily, RE causes high activity within these subtypes, which have high levels of recurrence (Fig 1F). Pairwise differences between FF, FBI, and RE KC activation patterns are measured by calculating the Hamming distance and difference in mean activation for each of the 250 sampled input patterns. While the feedback inhibition acts to decrease activity while slightly changing which neurons are active, RE induces larger changes in Hamming distance with the same change in mean activation (Fig 3H). Feedback inhibition and recurrent excitation are shown to each have unique roles, namely activation level decreases through FBI and changes in the underlying patterns of active neurons through RE.

**Fig 3.**
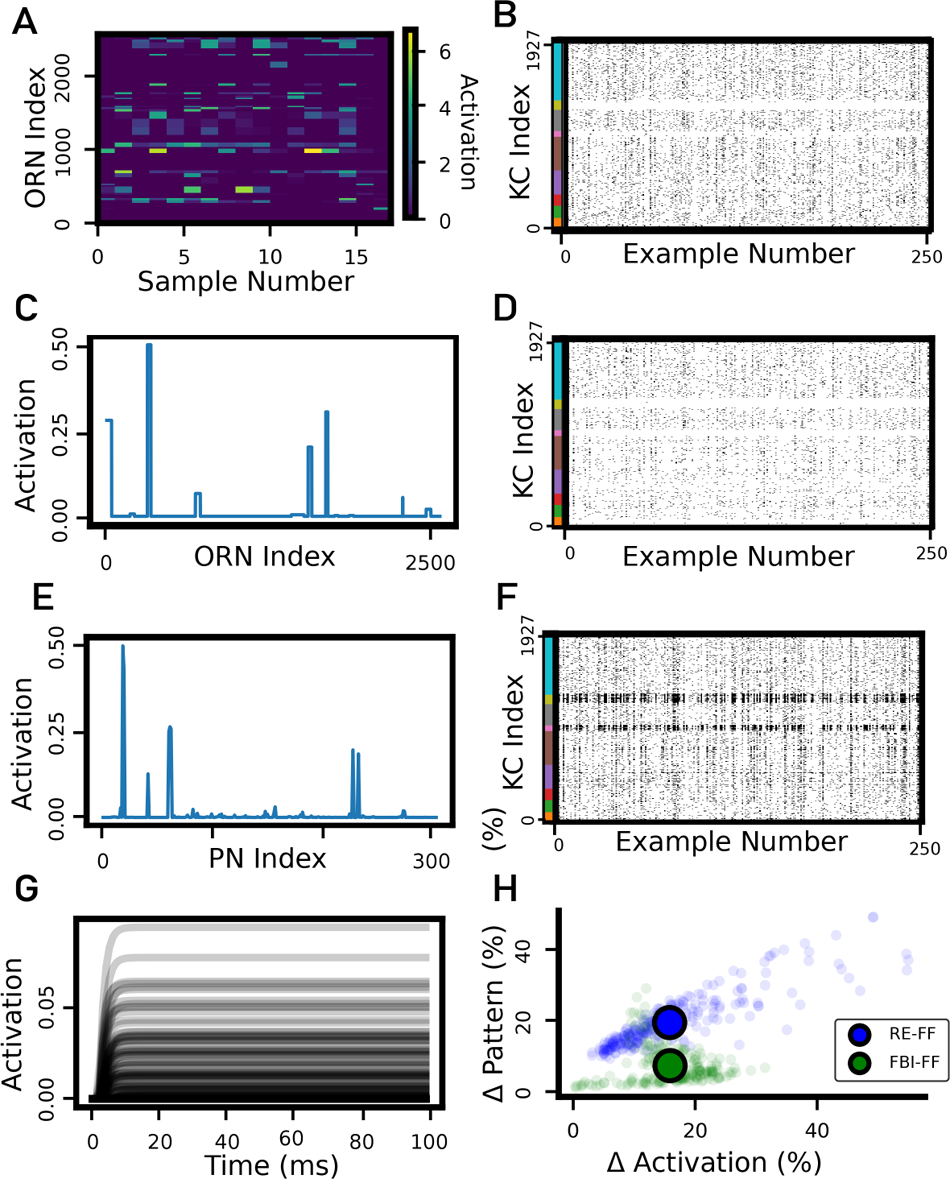
Simulated effects of feedback inhibition and recurrent excitation. (A) Experimentally recorded ORN activation patterns [45] to 17 different olfactory stimuli which were resampled as input to the network model (after being normalized by the standard deviation of the activation of all patterns). (C) Example resampled ORN activity used as network model input. Activity on the y-axis is defined as the activation divided by the max activation per neuron across all patterns. (E) Projection neuron activation with example ORN network model input. (G) KC activity traces corresponding to example ORN network model input with feedforward input only (i.e., no FBI or RE). Each line represents the activity of a different KC over time. (B) Binarized KC activity at the final time step for 250 different resampled ORN activation patterns with feedforward input only. (D,F) Binarized KC activity for resampled ORN activation patterns with FBI or RE respectively. For comparison, the levels of FBI and RE were chosen that decrease or increase the mean KC activation patterns by 15% respectively. (H) A comparison of the change in KC activation pattern (as measured by Hamming distance) vs. activation level (as measured by the absolute value of the mean activity difference) shows that when FBI and RE changed the activation level by the same mean amount relative to feedforward input only, RE caused a greater change in the activation pattern. The plotted Hamming difference is the Hamming distance as a percent of the total number of neurons. Individual smaller shaded markers represent measurements from each KC, while larger markers represent the mean across all KCs.

To further characterize the computational role of excitatory and inhibitory recurrence, we investigate how recurrent connectivity affects the dimensionality of KC activation patterns during sensory input as simulated for 100ms. The relative levels of recurrent excitation and feedback inhibition can be controlled in simulations and are measured with current ratios *C^RE^* and *C^F^ ^BI^* as the average ratio of recurrent excitation and feedback inhibition current to feedforward current respectively (as further defined in the Materials and methods section). During sensory input (Fig. 4A-B, 0-100ms), the activation level is different based on the level of RE. At low levels of RE (Fig. 4A), there is a minor recruitment of additional KC activation after initial sensory-triggered activation patterns. At higher levels of RE (Fig. 4B), there is a more pronounced recruitment of additional KC activation. Across 250 sampled input patterns at the end of the sensory input time period, some repeated features of KC activation patterns can be seen at low levels of RE (Fig. 4C), while more pronounced feature repetition in KC activation patterns can be seen at higher levels of RE (Fig. 4D). Effectively, more repeated features in KC activation patterns across sampled input patterns with higher RE corresponds to a decrease in the dimensionality of the KC activation patterns.

**Fig 4.**
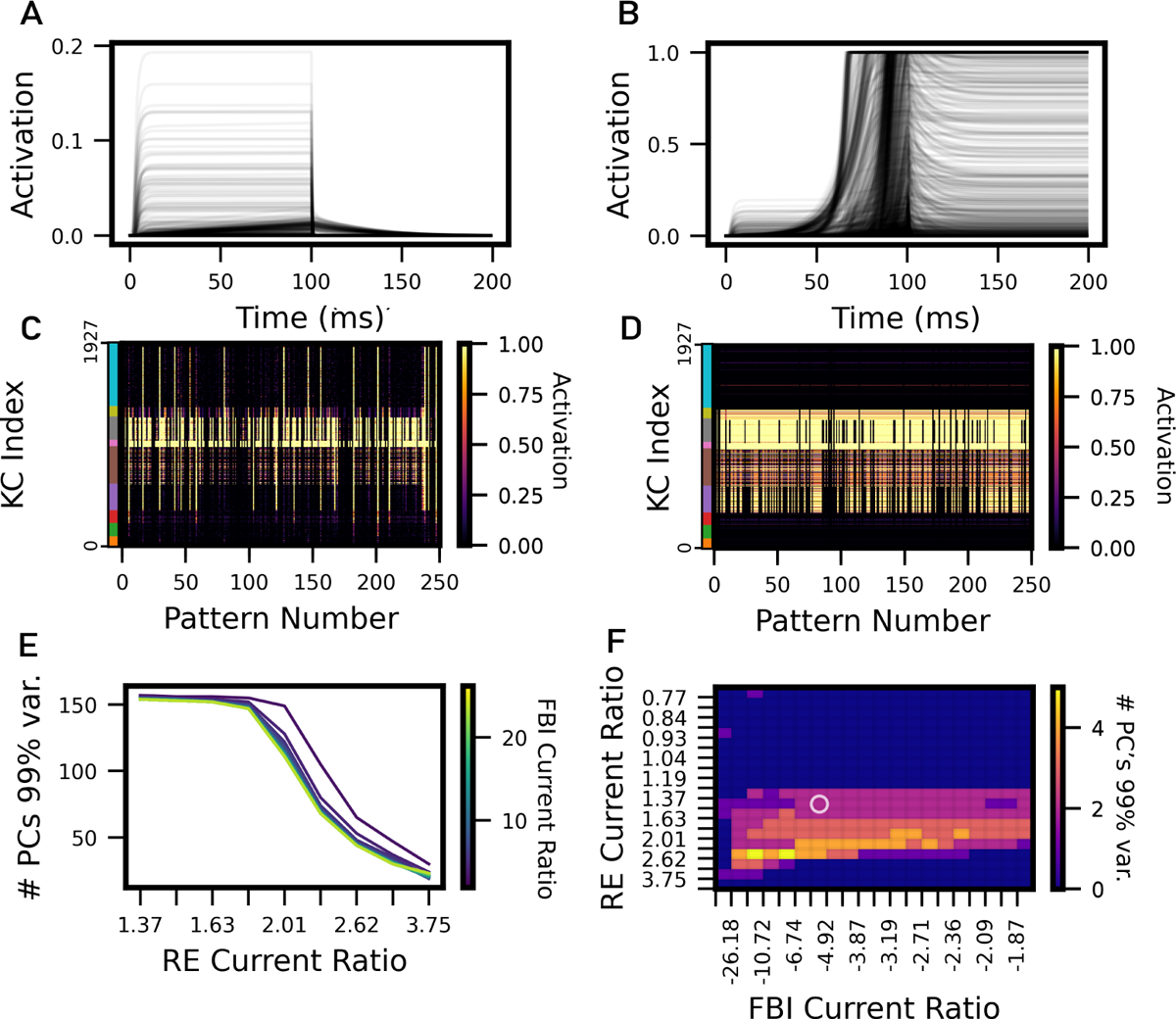
Recurrent dimensionality reduction and sustained activation. (A) The activity of each of the 1927 KCs across 200ms for an example pattern from a set of parameters with low enough RE that activation is not sustained after input was turned off. The feedforward input was turned off after 100ms. (B) In this case, an example is shown from a set of parameters where the RE is high enough to sustain activation after input was turned off. (C) Activity grid plot showing steady state activation of each KC across each of the 250 patterns at the right before input was turned off at 100ms. These patterns are from the same parameters shown in (A). (D) Activity grid plot showing steady state activation of each KC across each of the 250 patterns at 200ms. These patterns are from the same parameters shown in (B). (E-F) Characterization of dimensionality (measured as number of principal components to explain 99% of the variance in KC activity) as a function of RE and FBI level. (E) In the presence of sensory input (at 100ms), dimensionality is reduced with higher levels of RE. Higher levels of FBI also reduce dimensionality with a lower effect size. (F) In the absence of sensory input (at 200ms), intermediate levels of RE enable sustained activation patterns along multiple principle components. Note that at high and low levels of RE, activation patterns are either completed saturated or all zero respectively.

When dimensionality of KC activation is explicitly measured by the number of principle components needed to explain 99% of the variance, a monotonic relationship is observed where increased RE decreases dimensionality (Fig. 4E). The effect of FBI on KC dimensionality is less pronounced than RE, but it has the same effect direction, where increased FBI also causes decreased KC dimensionality. While the effect direction on dimensionality is the same for RE and FBI, the reasons are different: while RE recruits conserved activation patterns, FBI effectively suppresses smaller activation patterns.

We investigated how RE can mediate sustained activation patterns in the absence of input, while also reducing the dimensionality of the KC activation patterns as RE strength increases. To investigate sustained activation patterns, an initial 100ms sensory pattern presentation phase was followed by a 100ms response phase where there was no sensory input. At lower levels of RE, removing input during the response phase caused the activity of all neurons to decay to zero (Fig. 4A), but when RE was increased beyond a certain threshold, stable patterns were sustained after input was removed (Fig. 4B). As RE increased above this threshold, the dimensionality of these self-sustaining states generally increased as more KCs became active. As RE increased further, however, the dimensionality of these self-sustaining states decreased as KC activation saturated (Fig. 4F). The effect of FBI on self-sustaining states was less pronounced than RE, and was dependent on the level of RE. As RE increased, a higher level of FBI was necessary to maintain the dimensionality in self-sustaining KC activation and counteract saturation. The highest dimensionality sustained activation patterns were enabled at higher levels of FBI. Overall, after a sensory input, self-sustaining KC activation patterns with a compact set of repeated activation pattern components were driven by RE and modulated by FBI.

### Regime characterization

In addition to self-sustaining steady dynamics, additional qualitative properties are observed based on relative levels of RE and FBI (Fig. 5A). As RE increases, the resultant network activation patterns progress through characteristic properties, which we characterize as different regimes: 1)*Feedforward*, where the activation patterns are mostly unchanged from the input, 2) *Excitation*, where RE modifies the activation patterns, 3) *Self-sustaining*, where the activation patterns become self-sustaining, and 4) *Saturated*, where all activation patterns are fully saturated. When increasing FBI levels, we characterize two additional regimes: 5) *Stable FBI*, where activation approaches a pattern characteristic of FBI alone, and 6) *Oscillatory*, where there is a cyclic oscillation in the activation patterns. To quantify these different regimes, we introduce a series of metrics. A relative Hamming distance of greater than 2% of all KCs (39 neurons) is used to differentiate patterns as sufficiently distinct from the feedforward or FBI-only activation patterns. The *Self-sustaining* regime is defined as whether any KCs maintain activation after input is removed. Finally, the *Oscillatory* regime is defined if greater than 5% of patterns have oscillations (as detected by a decrease of 0.1% in the average activation pattern after reaching an initial maximum value). Given that the exact levels of RE and FBI are unknown, we introduce two criteria to characterize the biological plausibility of the identified regimes (Fig. 5B): 1) extreme levels of RE and/or FBI that are over an order magnitude larger than feedforward current are unlikely, and 2) the density of activation patterns should not exceed maximum observed levels of 15% [21]. Portions of the *Self-sustaining* and the entire *Saturated* regime have activation patterns that are denser than those observed in the fly, while the *Stable FBI* and *Oscillatory* regimes have FBI currents that are over an order of magnitude greater than the feedforward current. Portions of the *Feedforward*, *Excitation*, and *Self-sustaining* regimes match the introduced criteria, however, suggesting biological plausibility of these regimes.

**Fig 5.**
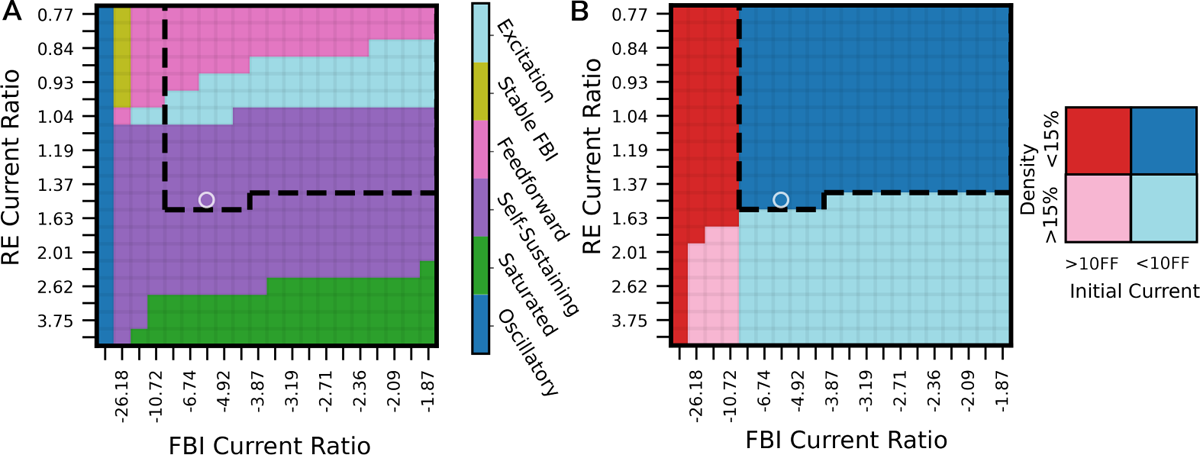
Activity regimes and biological plausibility. (A) Regimes of KC activation patterns were quantified as a function of relative levels of FBI and RE. (B) Biological plausibility of activation patterns based on criteria of 1) RE/FBI current within an order of magnitude of the feedforward current (blue vs. red), and 2) density of activation patterns within experimental observations (*<* 15%) (dark vs. light). Portions of the *Feedforward*, *Excitation*, and *Self-sustaining* regimes met these criteria for biological plausibility (dashed line). The relative levels of RE and FBI indicated with the white circle were utilized for more detailed analysis because they were representative of the parameters which sustained activation after input was disconnected. This parameter point was within the feasible region, and had the highest number of PCs to explain 99% of the variance after input is disconnected (see the corresponding to white circle in Fig. 4F).

### Stable State Analysis

In order to investigate the stability of the network dynamics around the identified self-sustaining point attractor states, we utilized a perturbation approach. Simulations utilized for the perturbation approach were conducted with four consecutive 100ms phases: 1) a resampled input pattern was presented, 2) the input pattern was removed, 3) a random Gaussian perturbation with the 0.1 times the mean of the input, and standard deviation equaling to that of the input data, was applied as input, and 4) all inputs were removed again. The KC patterns were then analyzed over time to identify whether the patterns return to their original input-driven self-sustaining states after perturbations were removed (Fig 6A). For this stability analysis, the parameters corresponding to the white circle in Fig. 5 from the regime analysis were used. These parameters yielded the highest number of principal components to explain 99% of the variance after input was turned off and also was within the feasible regime. The self-sustaining activation patterns were identified by evaluating the KC activation patterns that had stable non-zero activation patterns at t=100ms and t=200ms (872 activation patterns qualified out of a total of 2500 simulated input patterns). When analyzed after phase 2 in the perturbation simulation (after the original input patterns are removed at t=100ms), five discrete clusters were observed in KC activation patterns when projected onto the top two principal components of KC activity (Fig. 6B). KC activity returned to self-sustaining activation patterns after a perturbation was added and removed as shown in Fig. 6C, where the cosine distance to the KC patterns at t=200ms decreased during initial input presentation and removal, increased during perturbation, and decreased again following the removal of the perturbation. The KC activation patterns that settled into these five discrete clusters shared common activation patterns per KC subtype (Fig. 6D), where the activation patterns in the five discrete clusters mainly differed in which KC subtypes were active (Fig. 6E). The trajectories of KC activation patterns as they settled from input-driven states to self-sustaining states (Fig. 6F) predominantly demonstrated movement towards the closest of the five clusters (however, there was some overlap in the cluster four and five settling trajectories, likely due to the extinguishing of KCg-m activation in a subset of patterns that settled into cluster four). During perturbation, the activation patterns departed and returned to these discrete clusters (Fig. 6G). To quantify the input-driven KC activation dynamics in relation to the identified discrete clusters, the cosine similarity was measured between the KC activity and the closest cluster during the input-driven simulation phase. At the investigated RE level, the input-driven KC activation patterns approached these discrete clusters at varying speeds with an approximate final similarity of 0.9 (Fig. 6H). The input-driven KC activation dynamics was similarly quantified at varying levels of RE to investigate whether the identified discrete clusters were related to KC activation patterns across network assumptions with sparser KC activation patterns 6I). At lower levels of RE (corresponding to sparsity levels of 5-15%) the identified discrete clusters were still approached, however with lower final average similarities. Furthermore, a continuous relation was observed, where lower RE levels corresponded to lower final similarity values. Overall, point attractor dynamics were exhibited in perturbation simulations, including the following observations: 1) five discrete self-sustaining states were observed, 2) these discrete self-sustaining states were returned to following perturbations, and 3) the discrete self-sustaining states remained useful for describing input-driven KC activation patterns even under conditions with less RE and sparser KC activation patterns (and where these states were not even self-sustaining).

**Fig 6.**
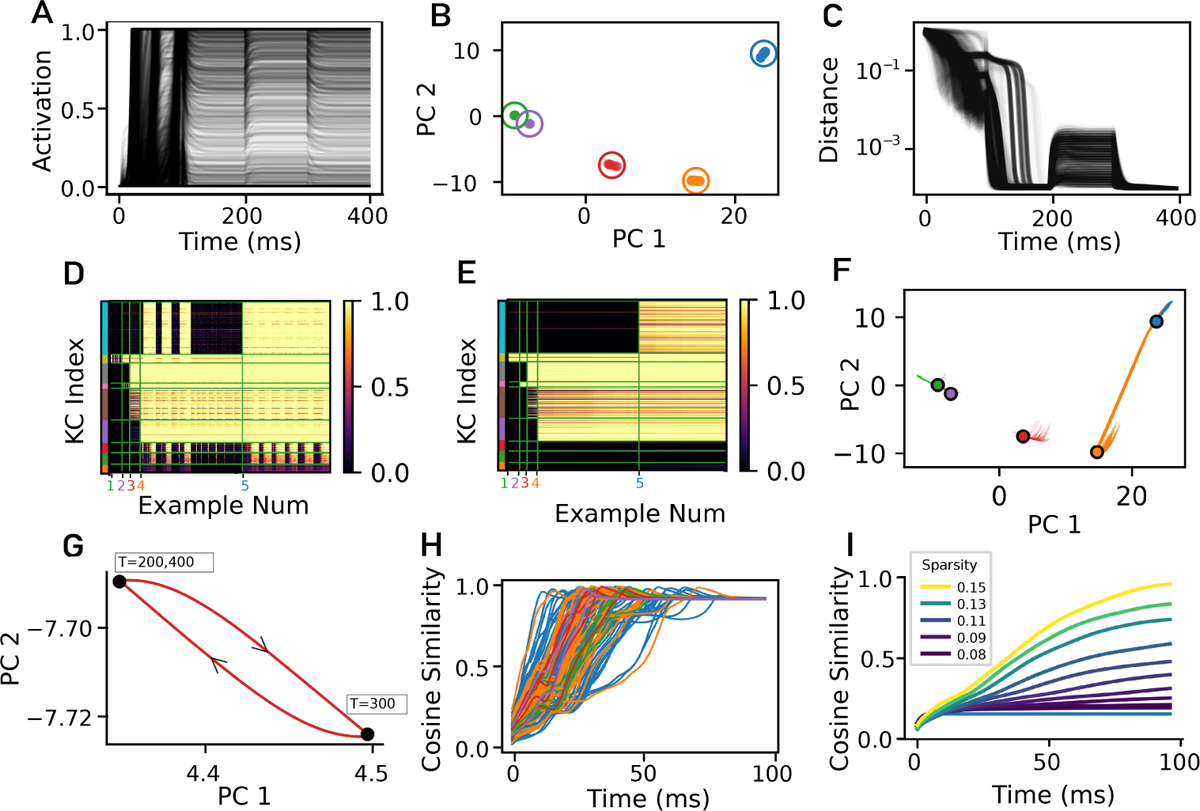
Point attractor dynamics quantified with a perturbation approach. (A) The activity of each of the 1927 KCs for an example input pattern during the four phases of simulated input: 1) re-sampled sensory input, 2) input removed, 3) perturbation input of a random shuffling of the input pattern, scaled by 0.1, and 4) input removed. Each phase lasted 100ms. (B) Five distinct clusters were observed in the activation along the top two principal components across input patterns at t=200ms. (C) An approach and return after perturbation to the network state at t=200ms was observed for the evaluated input patterns, as measured with the cosine distance of the KC activation over time to the activation at t=200ms (distance values are clipped to a minimum of 1e-4 for ease of visualization). (D,E) Activation patterns for all KCs at t=100ms and t=200ms respectively, where the x-axis is ordered per cluster as depicted in B and the y-axis is ordered per KC subtype as described in Fig. 1. Distinct clusters are composed of different combinations of active KC subtypes. (F) The trajectories of KC activation from t=100ms to t=200ms along the top two principal components for the evaluated input patterns show an approach to each cluster center. Each line corresponds to a distinct KC activation pattern and the points represents the average activation pattern for each of the clusters depicted in B. (G) For an example input pattern, there is a departure during perturbation (from t=200-300ms), followed by a return (from t=300-400ms) to the t=200ms pattern. (H) A visualization of the KC activation pattern approach to one of the cluster centers in B from t=0-100ms for each of the evaluated input patterns (as measured by cosine similarity, colored per the closest cluster at t=200ms). (I) KC activation pattern approach cluster centers in B across lower levels of RE. Color indicates sparsity as measured by the average number of neurons with activation greater than 0.01 divided by the total number of neurons. As in H, however all evaluated input pattern traces were averaged for each evaluated RE level (all lines in panel H average to the Sparsity=0.43 line in I). Note: For panels B-G, only patterns that were non-zero and stable at t=200 are depicted (872 out of 2500 where 1294 were zero and 428 were non-steady at t=200ms). For panels H-I, 250 input patterns were evaluated for each RE level investigated.

In order to investigate the behavioral implications of the self-sustaining point attractor states, the activation patterns of the MBONs (which have stereotyped roles in associating valences with sensory input as well as triggering approach and avoidance behaviors) were be analyzed for each state (Fig 7A). The behavioral roles for each MBON were delineated as positive, negative, or not described [5], so we therefore divided the MBONs into three corresponding behavioral sub-populations for further analysis. Note that all self-sustaining states triggered MBON activation that was distributed across all valence types (i.e., positive, negative, and not described) and there was no direct mapping between any self-sustaining state to a particular valence type (Fig 7 B). Across states, however, there were patterns of variation between which MBONs were active within each sub-population. The patterns of variation of each MBON sub-population were quantified by the correlation of each MBON sub-population across states (Fig 7 C-E). In the positive MBON sub-population, there were three distinct clusters of two states each, with variability within each cluster (Fig 7 C). In the negative MBON sub-population, there were two highly conserved clusters, one of two states and the other of four states (Fig 7 D). Each stable state had a unique distribution of contributing neurons from each of the valence types. The observed self-sustaining attractor states were also inspected from the perspective of sensory processing. The non-olfactory KCs (sub-types *γ*-d and *α*/*β*-p) were active in all self-sustaining attractor states and were the 1st and 2nd most active sub-types out of the 10 significant KC sub-types. The observed self-sustaining attractor states are not segregated across valences in the MBONs nor sensory-input type in the KCs; instead, more detailed patterns of variability were observed.

**Fig 7.**
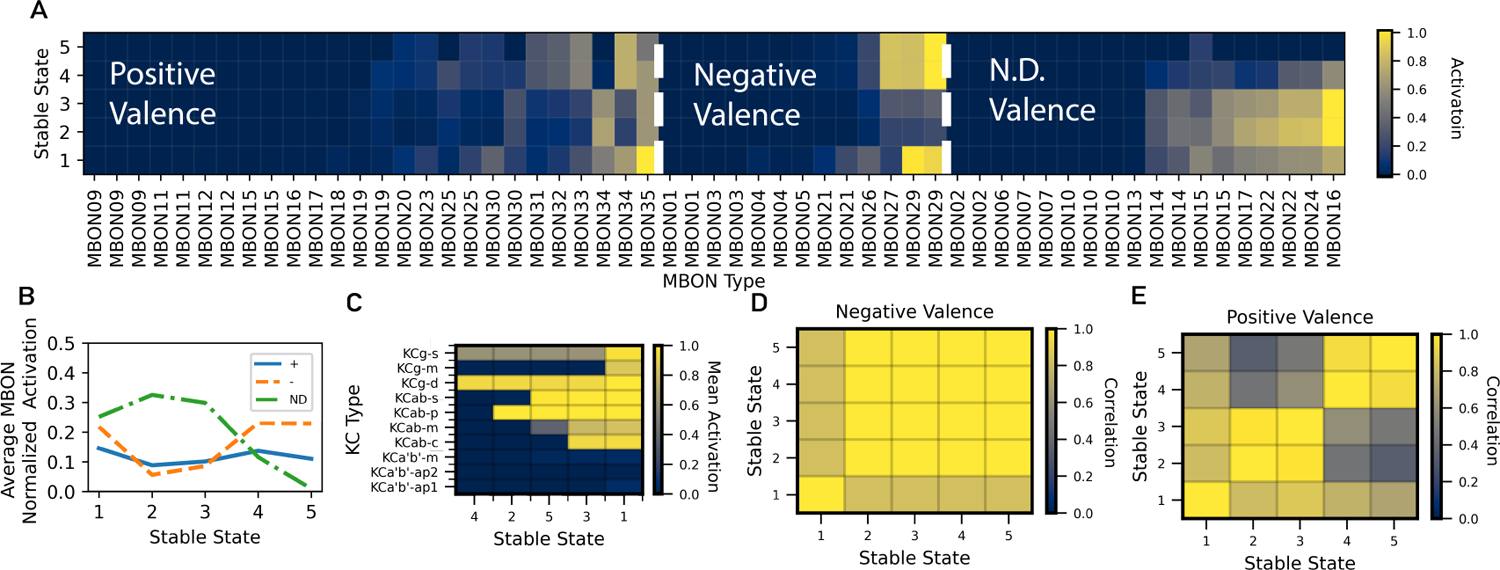
Behavioral properties of self-sustaining point attractor states. (A) The modeled MBON activation for each of the identified self-sustaining point attractor stable states is shown grouped by the valence that has been attributed to each MBON experimentally [5]. Activation was normalized by the maximum of each stable state, and MBON indices were sorted within each valence by the mean activation across all stable states. (B) For each stable state, the average MBON activation from A was compared per valence sub-population, showing a unique distribution across output neurons of positive, negative, and not described valence for each state. (C) For each stable state, the average KC activation per sub-type is shown with high activation in the KCg-d subtype for each state, and activation in the KCab-p for all but one state. The KCa’b’ neurons were not active in any state. (D-E) The patterns of MBON variation within the positive and negative sub-populations were compared via the correlation coefficient between activation patterns for each stable state. There were two clusters of negative valence activation with four out of the five states belonging to a single state. The positive patterns had three clusters of activation.

## Discussion

Neural networks with point attractor dynamics have rich theoretical computational properties [3], however it is largely unknown to what degree biological networks utilize point attractor dynamics and how the architecture of biological point attractor networks relate to their theoretical counterparts. In a series of analyses and network simulations, we investigated the role of recurrent connections among the predominant cell type (KCs) in the insect learning center (MB) for mediating dimensionality reduction and point attractor dynamics. For network simulations, synapse-level connectome data was utilized to create a model of the full input processing pathway from olfactory receptor neurons to Mushroom Body Output Neurons. From analyzing the network connectivity alone, it was observed that RE connections have high levels of symmetry and were balanced with feedforward input. In simulations, RE connections altered the overall network activation patterns and enhanced dimensionality reduction. The network dynamics investigated in simulations revealed a role for RE in enabling a small set of self-sustaining point attractor states. Additional dynamical network regimes were identified with varied levels of RE and FBI, and the more restricted biologically plausible sub-regimes encompassed these self-sustaining point attractor states. Behavioral implications of our analyses in the fly include the identification of a small set of MBON population components activated by these self-sustaining point attractor states as well as a role of predominantly non-olfactory KCs in processing olfactory sensory input.

Analyzing the recurrent KC network connectivity reveals properties of input integration and suggests a role of the recurrent connections in supporting stable attractor dynamics. For olfactory KCs, we found a correlation between the number of synapses from PN neurons carrying sensory input and the number of input synapses from other KC neurons, suggesting that balanced feedforward and recurrent excitation input is an invariant feature across the MB. Furthermore, while the numbers of synapses in PN and APL are correlated [1], the correlation between PN and KC recurrent neuron inputs is larger (r=0.63 versus r=0.41 for PN and APL synapses), suggesting the relative importance of balanced feedforward sensory input and recurrent excitation versus balanced feedforward input and FBI. We also analyzed the recurrent connectivity of the KCs by calculating symmetry to gain insight into the role of these connections to support stable attractor dynamics. A fully symmetric weight matrix in a recurrent neural network implies stable dynamics by guaranteeing the existence of a Lyapunov function [12, 25] Computationally, these stable network dynamics can be used for discrete pattern retrieval [22] and for quadratic optimization [23]. Although a symmetric weight matrix is a sufficient condition for dynamical stability, this condition is not necessary. For example, random damages to the connections of a Hopfield network destroy the symmetry of the weight matrix, but the attractor dynamics typically remain intact [3]. Our finding that the recurrent excitation (mediated through KC-KC) and FBI (mediated through KC-APL-KC) have high levels of symmetry justifies further investigation into the role of these recurrent connections in enabling stable attractor dynamics.

We performed network simulations to investigate the effects of recurrent connections on KC neuron activation patterns and found that while FBI changes the overall network activation level, RE changes the underlying activation patterns. This role for FBI of suppressing KC activation and enabling sparse activation patterns is consistent with previous findings for the APL [33, 39] and for inhibitory neurons more broadly [49]. In contrast, RE in our simulations changed the underlying activation patterns (as measured with a larger Hamming distance in network activation when adding RE vs. FBI). The role of FBI as providing general inhibition can be explained by the presence of a single inhibitory neuron per hemisphere, the APL neuron, whereas the effective FBI weight matrix is the outer product of the KC*→*APL and APL*→*KC weights. The KC*→*APL weights determine how the overall activity level of the KC population is encoded in the APL activation, and the APL*→*KC weights determine how inhibition is distributed among the population of KC neurons. In contrast, the RE weight matrix is full rank and causes a larger modification in the structure of the KC activation patterns than FBI.

The information processing in the MB without KC RE has been well studied theoretically, where sensory features in the PNs project randomly [9] to a much larger population of KC neurons (which have sparse activity enforced by APL FBI), and in turn have learned weights to a compact set of MBONs [20]. High dimensionality is typically seen as a key feature in KC activation [34] in the expansion architecture between PNs and MBONs. As has been observed in simulations of the larval MB [16], we show here that KC RE caused a pronounced dimensionality reduction in our simulations of the adult MB, which differs notably from the larval MB by having a larger ratio of KC neurons to PN neurons. Furthermore, when we varied the relative strength of RE in simulation (since this quantity has not been observed experimentally), we found that there was a rapid phase transition to a large level of dimensionality reduction. While such dimensionality reduction might seem to counter the computational benefits of a high dimensional expansion in KCs, the feedforward sensory inputs from PNs are still projected and mixed into a larger population of KCs. The dimensionality reduction by RE in KCs over time in a layer before MBONs, therefore, can be interpreted as an additional computational transformation in the processing of the MB. The role of recurrent connectivity in dimensionality reduction is analogous to reduced dimensionality seen with high levels of recurrent connectivity experimentally in other neural circuits critical to memory formation, such as the hippocampal CA3 [30], [31], and in cortical areas such as those involved in decision making and motor control [17], [19], [26].

Properties of self-sustaining point attractor dynamics were observed in our simulations with recurrent connectivity, whereby, after presentation of a sampled sensory pattern, a discrete set of stable activation patterns persisted in the absence of external input. Furthermore, after applying a perturbation to these activation patterns, the activity in the network returned to the original stable activation pattern. While point attractor dynamics have been observed experimentally [18], [26], [29], [40], [46], [47], [51], several fundamental questions remain as to the relationship between theoretical point attractor networks [3], and those in biological neural networks. While in theoretical models, the number of possible point attractor states cannot exceed the number of neurons in the recurrent neural population [3], [43], it is difficult to probe such a high-dimensional attractor landscape experimentally. *In vivo* studies have necessarily focused on studying the dynamics of a number of attractor states far less than the total number of neurons in a population [26]. In our simulations, where we use a sampling approach to probe the attractor landscape, we observed a maximum of up to five self-sustaining attractor states in the population of 1927 modeled KCs. This relatively small number of point attractor states suggests that each observed state likely would not be linked to unique low-level single sensory experiences such as specific odors, but would rather be useful downstream for higher-level categorical representations.

While our sampling approach is not exhaustive over all possible sensory input patterns, and thus possibly misses smaller basins of attractions, the point attractor states observed in simulations were consistent across samples. Sources of additional network input such as additional MB cell types not included in our network models (e.g. dopaminergic neurons) or neurons in other neuropils will undoubtedly affect neural activation dynamics. Nevertheless, network simulations support a role for the recurrent KC connectivity along with feedback inhibition of the APL to support self-sustaining point attractor dynamics.

In addition to self-sustaining point attractor dynamics, there are other possible dynamical regimes observed when varying the level of RE and FBI. We quantified these network dynamics into six characteristic regimes and assessed their biological plausibility. At the extremes, high levels of RE led to complete saturation, and high levels of FBI led to oscillatory activation. During oscillatory activation, each cycle consisted of strong FBI that reduced all KC membrane activation to sub-threshold levels, followed by deactivation of the inhibitory APL neuron, followed by the reactivation of KCs. At low levels of RE, network dynamics either approximated feedforward only input (with low FBI) or approximated the same output as FBI only (at higher levels of FBI), regimes which we refer to as “Feedforward” and “Stable FBI” respectively. At intermediate levels of RE and FBI, however, two new network dynamic regimes were observed: 1) “Excitation,” where activation patterns are changed from the feedforward or FBI only case, and 2) the “Self-sustaining” dynamics as described previously. To assess the biological plausibility of these regimes in relation to experimental results, we propose two criteria: 1) maximum sparsity levels of KC activation comparable with experimental observations (as high as 15% [21]) and 2) relative levels of FBI and RE current within one order of magnitude as feedforward input current. According to these criteria, portions of the the “Excitation”, “Feedforward”, and “Self-sustaining” regimes are biologically plausible. In the “Excitation” regime, which had sparser overall levels of activation than the “Self-sustaining” regime, the KC activation patterns still approached one of the identified self-sustaining point attractor states. This suggests that recurrent connections bias activity towards the identified self-sustaining point attractor states, even in the cases where RE is not strong enough to support these activation patterns without any input.

We also investigated the functional implications of the identified self-sustaining point attractor states in the context of both processed sensory inputs to the MB and behavior-influencing outputs from the MB mediated through MBONs. The minority of KCs integrate non-olfactory sensory input (8% of all KCs are in KCg-d, KCab-p). Previous studies [16] have raised the possibility that these non-olfactory KC’s may play an outsized role in influencing certain MBON activation, due to their relative over-representation in the total number of overall KC-MBON synapses. From analyzing network connectivity, it is apparent that these non-olfactory KCs had recurrent connectivity to olfactory KCs, indicating a role for non-olfactory KCs in processing olfactory input. In network simulations, the presence of these non-olfactory KCs in all self-sustaining point attractor states suggests a further role of non-olfactory KCs influencing overall network activation dynamics. One way to investigate the behavioral role of the KC self-sustaining point attractor states is in the context of individual MBON activation, which have been stereotyped by their role in associating positive and negative valences to sensory input, as well as triggering approach and avoidance behaviors [38]. When investigating the evoked MBON activation patterns for each self-sustaining point attractor state, we found that there was no clear division between states preferentially recruiting positive or negative valenced MBONs. This indicated that the stable states may not have been directly associated with influencing approach and avoid decision-making, but could influence additional perceptual dimensions (as opposed to behavioral dimensions). Furthermore, self-sustaining states were clustered into different sets of components among the positive and negative valence compartments in the MB, potentially motivating future investigations (especially in the context of the multi-layered logic of MBONs [5]). Overall, self-sustaining point attractor states lead to testable predictions in the fly that: 1) non-olfactory sensory input influences all MBON-learned behaviors (not just the MBONs that receive input from non-olfactory KCs) and 2) Recurrent KC connections mediate a small set of components in MBON activation (that are not organized by positive or negative valence).

While many simplifications in our computational approach are necessary, they may limit the applicability of our conclusions and can be investigated in follow-up work. In the network architecture, we focused on the role of KC recurrent connections, and thus did not include all MB neuron types in our model, such as dopaminergic and dorsal paired medial neurons, which could be incorporated into future work. An additional simplifying assumption utilized is that all KC-KC connections are excitatory. Recent experimental results suggest that convergent KC-KC connections, particularly in the gamma lobe may participate an inhibitory neuromodulation [35]. While we filter out convergent KC-KC connections and include non-lobe calyx connections in our analysis, our work motivates further quantification of neruotransmission at non-convergent KC-connections to inform future modeling and the interpretation of our results. For input to our network, we utilized olfactory patterns considering that odor patterns are important for driving many of the fly’s behaviors, and have been the most studied sensory modality in the MB [20], however including additional sensory modality inputs would give a more complete description. Our simulations were targeted at capturing fundamental architecture-based and firing rate-based computational properties and thus we did not explicitly model spiking neuron activity nor complex time-varying sensory input patterns [7]. In addition, the morphologies of individual neurons were not explicitly modeled and phenomena such as localized inhibition [2] were not included, which may further influence network dynamics. A model detail that could be better informed by future experimental studies and which could affect network dynamics is the threshold of KCs which may vary from cell to cell. For our analysis, we constrain KC thresholds as in previous MB models [16] to support sparse KC activation. Additionally, the weight matrix values in the network simulations were derived from synapse counts, which is an indirect measure of connectivity strength, but one that has been shown to accurately predict total synaptic surface area [6].

## Conclusion

These analyses and simulations of the complete synapse-level recurrent connectivity of the predominant cell type in the insect MB yield new insights into the computational principles of this circuit, especially by (1) quantifying its enhanced dimensionality reduction process and (2) identifying the presence of a compact set of self-sustaining point attractor states. Since recurrent networks are ubiquitous across organisms, the observed relationship between the recurrent connectivity in the MB and the properties of the point attractor dynamics described here might serve as a base model to be compared with circuits in higher-level organisms, potentially enabling the identification of computational principles governing the generic modes of operation for recurrent neural populations.

## Materials and methods

### Network Structure

The effective connectivity weights in the connectome-constrained network model (*_W_ORN →PN* _, *W*_*PN →KC* _, *W*_*KC→APL*, *_W_APL→KC* _, *W*_*KC→KC* _, and *W*_*KC→MBON*) were constructed by utilizing the count of synapses between individual neurons of the ORN, PN, KC, APL, and MBON neuron types in a recent publicly released connectome dataset [44], accessed using https://neuprint.janelia.org/. To filter out potentially spurious connections and constrain analysis to the mushroom body, only connections with greater than five synapses located within the mushroom body were included.

Feedback connections of PN*→*ORN and MBON*→*KC, were omitted from the model due to a diminished synapse count relative to the number of feedforward synapses (1.3% and 1.5% respectively). In the network model, reciprocal connections between KCs at convergent/rosette synapses were not included because it is unknown whether activation between KCs can be propagated at these synapses [48]. These convergent/rosette synapses are annotated [44], however, and represent 75% of all annotated recurrent connections. The reciprocal connections between KCs at convergent synapses were identified for removal if they exist at a synaptic site containing any cell type other than a KC neuron.

### Biologically-constrained network input

Network simulations were repeatedly performed with *N* re-sampled ORN response patterns drawn from the distribution of reported experimentally recorded activation levels [45]. This dataset includes the response of 31 different glomeruli to 17 different olfactory stimuli. Each re-sampled ORN response pattern, *r_n_*, includes multiple (17-142) neurons from the same glomeruli, which share an activation level sampled from the 17 different olfactory stimuli responses. The set of PN activation patterns for downstream simulation were defined as *X* = *RW_ORN→P N_ /σ_P N_* (Fig 3), where *R* was the row-wise concatenation of all *r_n_*resampled ORN reponse patterns and *σ_P N_* was a normalization factor calculated as the standard deviation of all elements in *RW_ORN→P N_*.

### Leaky integrator neuron model

The dynamics of the KCs and APL neuron were calculated as leaky integrator neurons, where the membrane potential, *u*(*t*), was updated by integrating the input current *j*(*t*):

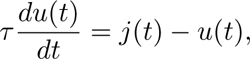

where the time constant *τ* was modeled as 20ms of KCs and 10ms for APL neurons.

The activation level of each neuron was modeled as *g*(*u*(*t*) *− θ*) by including a neuron-specific threshold, *θ*, and a nonlinear sigmoidal function, *g*(.), which squashed the activation level between 0 and 1. The activation of all KC neurons, *h*(*t*) = *g*(*u_KC_ − θ_KC_*) and APL neuron, *z*(*t*) = *g*(*u_AP L_ − θ_AP L_*) were calculated from the vectors of membrane potentials (*u_KC_, u_AP L_*) and thresholds (*θ_KC_, θ_AP L_*) for KC and APL neurons, respectively.

### Simulation of network variants

The simulated input current to KC neurons is a combination of feedforward olfactory input, recurrent excitation, and feedback inhibition. The total current delivered to the *i*th KC in each time step is modeled as:

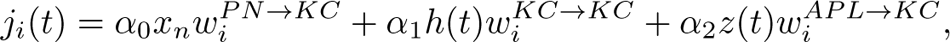

where *x_n_* is the PN activation vector for a single input pattern and *w_i_* denotes the column of each set of connectivity weights corresponding to the *i*th KC. The scalars *α*_0_, *α*_1_, and *α*_2_, control the relative strength of the feedforward input, recurrent excitation, and feedback inhibition.

To scale the feedforward input relative to the activation function, a scaling factor was calculated to scale the maximum input current across all KCs to one for a single input pattern, *α^n^* = 1*/*max(*x_n_W_P N→KC_*). For uniformity of simulation across all input patterns, the feedforward scaling factor *α*_0_ was calculated as the median value of all *α^n^*. The scaling factors *α*_1_ and *α*_2_ were adjusted for different network variants to investigate the properties of the network under various levels of recurrent excitation and feedback inhibition.

The threshold of each KC, *θ_KC,i_*, was constrained so that each KC responds to 5% of the input patterns when driven by feedforward input alone. When driven with a constant input pattern, the steady state membrane potential value is equal to the input current (following the solution to the first order linear non-homogenous differential equation). Thus, *θ_KC,i_*was set to the 95th percentile value of *α*_0_*Xw^P^ ^N→KC^*. An intermediate value of 0.5 was used for the single *θ_AP L_* in the simulation.

The current delivered to the APL neuron in each time step was defined as:

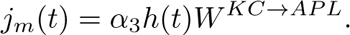

Similarly, to scale the APL input relative to the activation function, the scaling factor *α*_3_ was calculated to scale the APL input current to a median value of one across input patterns, *α*_3_ = median(1*/* max(*h_ss,n_W_KC→AP L_*)), where *h_ss,n_* is the KC activation level at the feedforward input-only steady state solution.

To provide interpretable measures of the relative recurrent excitation and feedback inhibition, current ratio terms *C^RE^* and *C^F^ ^BI^* were calculated as a ratio of the recurrent excitation and feedback inhibition current to feedforward current, respectively. For a single input pattern and a single KC neuron, the recurrent excitation ratio term was computed as:

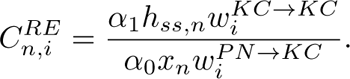

Similarly, the feedback inhibition ratio term was computed as:

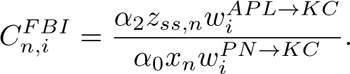

For these calculations, the KC and APL activation levels were set to their feedforward input-only steady state solutions, (*h_ss,n_* and *z_ss,n_* respectively). The current ratio terms, *C^RE^* and *C^F^ ^BI^*, were calculated as the average across all KC neurons and input patterns.

## Acknowledgments

Research reported in this publication was supported by DARPA Grant HR00111990038 and an internal grant from the Johns Hopkins University Applied Physics Laboratory.

## Disclaimer

This material is based on work while one of the authors (GMH) was supported by (while serving at) the National Science Foundation. Any opinions, findings, conclusions, or recommendations expressed in this material are those of the authors and do not necessarily reflect the views of the National Science Foundation. This article was prepared while Dr. Grace M. Hwang was employed at Johns Hopkins University and the National Science Foundation. The opinions expressed in this article are the author’s own and do not reflect the view of the National Institutes of Health, the Department of Health and Human Services, or the United States Government.

